# Quantitative proteomics reveals extensive lysine ubiquitination in the Arabidopsis root proteome and uncovers novel transcription factor stability states

**DOI:** 10.1101/2021.01.07.425780

**Authors:** Gaoyuan Song, Damilola Olatunji, Christian Montes, Natalie M Clark, Yunting Pu, Dior R Kelley, Justin W Walley

## Abstract

Protein activity, abundance, and stability can be regulated by posttranslational modification including ubiquitination. Ubiquitination is conserved among eukaryotes and plays a central role in modulating cellular function and yet we lack comprehensive catalogs of proteins that are modified by ubiquitin in plants. In this study, we describe an antibody-based approach to enrich peptides containing the di-glycine (diGly) remnant of ubiquitin and coupled that with isobaric labeling to enable quantification, from up to 16-multiplexed samples, for plant tissues. Collectively, we identified 7,130 diGly-modified lysine residues sites arising from 3,178 proteins in Arabidopsis primary roots. These data include ubiquitin proteasome dependent ubiquitination events as well as ubiquitination events associated with auxin treatment. Gene Ontology analysis indicated that ubiquitinated proteins are associated with numerous biological processes including hormone signaling, plant defense, protein homeostasis, and root morphogenesis. We determined the ubiquitinated lysine residues that directly regulate the stability of the transcription factors CRYPTOCHROME-INTERACTING BASIC-HELIX-LOOP-HELIX 1 (CIB1), CIB1 LIKE PROTEIN 2 (CIL2), and SENSITIVE TO PROTON RHIZOTOXICITY (STOP1) using site directed mutagenesis and *in vivo* degradation assays. These comprehensive site-level ubiquitinome profiles provide a wealth of data for future studies related to modulation of biological processes mediated by this posttranslational modification in plants.

## Introduction

Ubiquitin is a well-established posttranslational protein modification (PTM) that impacts nearly all aspects of plant biology (51). Covalent attachment of ubiquitin to substrate proteins occurs in a step-wise fashion involving E1 (ubiquitin activating), E2 (ubiquitin conjugating), and E3 (ubiquitin ligase) enzymes (27). The attachment of ubiquitin to protein substrates can result in many functional outcomes, including protein degradation or changes in subcellular localization. While ubiquitin is typically attached to lysine residues, it can also be covalently linked to cysteine, serine, threonine and the amino terminus of target proteins (1, 6, 9, 16, 28). In addition to positional complexity there is also oligomeric complexity whereby ubiquitin attachments to substrates can occur in various numbers of monomers (37). In Arabidopsis, over 1,500 annotated genes are linked to ubiquitin pathways suggesting that the biological processes involving this PTM are extensive (50). While ubiquitin 26S proteasome (UPS) mediated protein degradation has been demonstrated for most plant hormones (24), very little is known about the corresponding ubiquitin attachment(s) underlying such regulated proteolysis.

Given the importance of ubiquitin in modulating protein function a range of approaches have been used to identify ubiquitinated proteins in plants. One approach has been to use ubiquitin-associated domains or ubiquitin interaction motifs to affinity purify proteins with ubiquitin conjugates (22, 32, 33). Saracco et al. (40) developed a transgenic Arabidopsis line containing a 6xHis-UBQ tagged ubiquitin line, which has been used in combination with an additional enrichment step based on the HHR23A ubiquitin binding region (40) or tandem ubiquitin binding entities (2, 25) to purify ubiquitinated proteins. These methods are powerful for identifying ubiquitinated proteins but are unable to comprehensively identify the exact amino acid attached to ubiquitin because the enrichment occurs at the protein level. In other eukaryotic systems, the gold standard approach uses antibodies that recognize the di-glycine (diGly) remnant of ubiquitin and the ubiquitin-like protein Related-to-Ub-1 (RUB1/NEDD8), which remains following trypsin digestion, to enrich ubiquitin modified peptides (14, 26). This diGly antibody enrichment approach has been used on several plant species, yet the number of identified ubiquitin sites lags behind non-plant studies (7, 20, 30, 56, 58, 60). Thus, a combined fractional diagonal chromatography (COFRADIC) approach was developed to identify exact ubiquitin sites and expand our known repertoire of ubiquitinated Arabidopsis proteins (54). While COFRADIC has enabled the identification of the largest number of ubiquitin sites (3,009) in plants it requires complex *in vitro* chemical modification and sample processing steps, making it difficult to carry out (Supplemental Table 1).

Here, we report a diGly based method, using commercially available antibodies, for quantitative profiling of ubiquitinomes in plants. Our approach utilizes an isobaric tag-based mass spectrometry labeling method that enables sample multiplexing and relative quantification of changes in ubiquitin levels at specific amino acid sites. Isobaric labeling has an advantage over label-free proteomics approaches in that it does not suffer from missing values between samples, which is a challenge for large-scale studies. Using these methods, we report an extensive catalog of ubiquitin attachment sites from Arabidopsis root tissue. The identified ubiquitination sites occur on proteins from diverse functional categories and include many well-known 26S ubiquitin proteasome substrates for which the specific modified site was unknown. The ubiquitination sites identified in this study provide a rich resource for future mechanistic studies investigating protein function and deepens our understanding of the Arabidopsis root proteome.

## Results and Discussion

### Detection and quantification of ubiquitin-modified proteins in Arabidopsis roots

Despite widespread interest in protein ubiquitination, the ability to robustly carry out ubiquitinome analyses remains a challenge in plants. Thus, we sought to streamline the workflow and generate a reproducible protocol using commercially available reagents to increase utility for the community and expand the catalog of known ubiquitination sites on plant proteins (Figure 1A). We selected Arabidopsis roots because they serve as a key organ for investigating regulation of gene expression; both hormone signaling and protein turnover via the 26S proteasome have been implicated in regulating root growth and development (39, 52). We initially tested a small-scale enrichment from 2 mg of total peptides and 40 μl of pan anti-diGly remanent antibody conjugated to agarose beads (PTM Biolabs). The recovered peptides were analyzed using a 150 min 1-dimensonal reverse phase liquid chromatography gradient to deliver the peptides for tandem mass spectrometry (1D-LC-MS/MS). From this initial test experiment, we were able to identify 1,178 diGly-modified lysine residues (Supplemental Table 2). Encouraged by this initial result, we scaled-up to enrich from 20 mg of peptides prepared from roots treated with 100 μM of the 26S proteosome inhibitor bortezomib (BTZ) using 400 μl of anti-diGly conjugated beads. From this enrichment, we recovered sufficient peptides to further fractionate the samples and perform 2D-LC-MS/MS. This large-scale label-free analysis resulted in the identification of 3,955 diGly-modified lysine residues from a single sample (Figure 1C and Supplemental Table 2), which represents a sizeable increase in the depth of ubiquitinome profiling in plants.

**Fig. 1.**
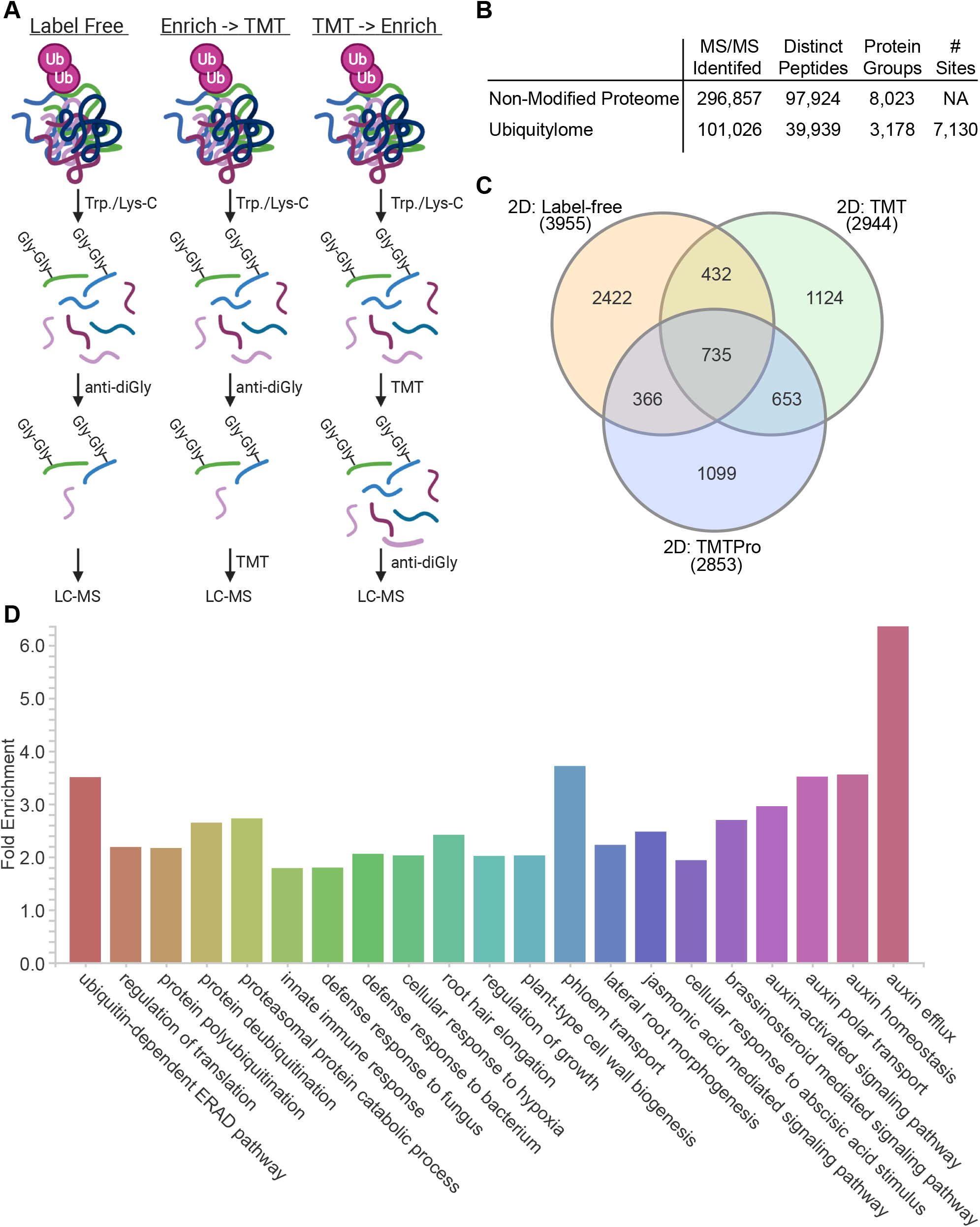
Quantitative proteomic analysis of the Arabidopsis root ubiquitinome. (A) Schematic of workflows tested for profiling diGly modified lysine residues. (B) Summary of identified spectra, peptides, proteins (i.e. protein groups), and diGly sites. NA, not applicable. (C) Overlap in the diGly sites detected using the Label Free and Enrich -> TMT workflows. The Enrich -> TMT approach was tested using both TMT and TMTpro reagents. (D) Selected Biological Process GO terms that are enriched (FDR < 0.05) among the diGly modified proteins. Supplemental Table 1 contains the full list of enriched GO terms.

While label-free approaches have been useful for identifying diGly sites, isobaric chemical tags for sample multiplexing offer several advantages for quantifying post-translational modifications (PTMs). Isobaric tags enable comparisons of up to 16 samples in a single mass spectrometry analysis. As a result, isobaric tag multiplexing improves throughput and significantly minimizes the number of missing peptides quantified across all experimental conditions, which is a major drawback of label-free analyses. Furthermore, multiplexing facilitates a decrease in the starting material needed, from each sample, for enrichment. Thus, we tested the compatibility of isobaric labeling using tandem mass tags (TMT) with diGly enrichment. In order to investigate the influence of hormone treatment and ubiquitin proteasome dependent modifications, we performed separate exogenous treatments with the phytohormone indole-3-acetic acid (IAA) (19) and proteasome inhibitor bortezomib (BTZ) (18). For these experiments, we used three biological replicates of mock (DMSO), 1 μM IAA, or 100 μM BTZ treated 10-day old Col-0 roots. First, we explored TMT labeling (2 mg peptides per sample; 18 mg total peptides) followed by diGly enrichment (“TMT -> Enrich” in Figure 1A). However, this method only recovered 10 diGly sites (data not shown). This is likely due to TMT labeling the primary amine of the diGly remnant, thereby inhibiting enrichment using the diGly antibody, as has been suggested (38, 48).

Next, we tested an alternative approach where we enriched with anti-diGly from 2 mg of peptide/sample and then TMT labeled the immunoprecipitated peptides (“Enrich -> TMT” in Figure 1A). Excitingly, this resulted in quantification of 2,944 diGly-modified lysine residues (Figure 1C and Supplemental Table 2). The original TMT labels used here allowed for multiplexing of up to 11 samples, but a newer TMTpro version was recently developed that can multiplex up to 16 samples. Additionally, TMTpro labels have altered chemical properties and are more hydrophobic compared to TMT 11-plex. Thus, we also tested the newer TMTpro labels for compatibility using an aliquot of the same peptides (2 mg each sample) from the original TMT analysis for diGly enrichment. Critically, we were able to quantify a similar (2,853) number of diGly sites using TMTpro multiplexing (Figure 1C and Supplemental Table 2). Furthermore, the “Enrich -> TMT” method was highly reproducible with average Pearson correlation values between biological replicates of 0.972 and 0.995 for the TMT and TMTpro analyses, respectively. Together, these results demonstrate the ability to analyze the plant ubiquitinome using reagents that enable multiplexing of up to 16 samples per mass spectrometry analysis, which facilitates large-scale quantitative studies.

To gain insight into the composition of the ubiquitinome we first merged the results of the experiments described above. In total, we identified 7,130 diGly-modified lysine residues sites arising from 3,178 proteins (Figure 1B and Supplemental Table 2), which represents a notable increase in the coverage of the Arabidopsis ubiquitinome (Supplemental Table 1). We used PANTHER to determine enrichment of Gene Ontology (GO) categories among these proteins (34). This analysis uncovered 553 enriched GO terms spanning a wide range of biological processes including (de)ubiquitination, translation, cell wall biogenesis, transport, and lipid metabolism (Figure 1D and Supplemental Table 3). Additionally, GO terms associated with plant development, growth, and defense were enriched. As a final example, several terms associated with phytohormone pathways including abscisic acid, auxin, brassinosteroid and jasmonic acid were observed. This analysis highlights the extensive biological functions which are potentially impacted by ubiquitination and is consistent with the UPS playing a key role in nearly all aspects of plant biology (51).

### Identification of potential UPS substrates

The attachment of ubiquitin to proteins is known to result in many different functional outcomes including altering protein stability by the UPS. Thus, to identify potential UPS substrates we examined quantitative changes in protein abundance and diGly-modified lysine levels following treatment with 100 μM BTZ, assuming that UPS substrates would increase in both abundance and ubiquitination following BTZ treatment. From this analysis, we found that 1,336 diGly sites on 691 proteins accumulate following BTZ treatment (Supplemental Table 2; Fold-change (FC) >1.1 q-value <0.1). To complement this finding, we also quantified protein abundance by TMTPro labeling an aliquot of the peptides that were used for diGly enrichment and analyzed them by 2D-LC-MS/MS. From this we identified 213 proteins that increased in abundance following BTZ treatment as potential UPS substrates (Figure 2A Supplemental Table 4; FC >1.1 p-value < 0.05). To add further evidence and create a high-confidence list of UPS substrates we compared these data with the list of diGly modified proteins and found evidence for 104 proteins as being both diGly modified and exhibiting an increase in protein level following BTZ treatment (Figure 2A). We examined the PANTHER protein class annotations for these 104 high-confidence UPS candidate proteins and found they are comprised of proteins involved in processes in including transcription and translation, defense, cytoskeleton, and transport (Figure 2B).

**Fig. 2.**
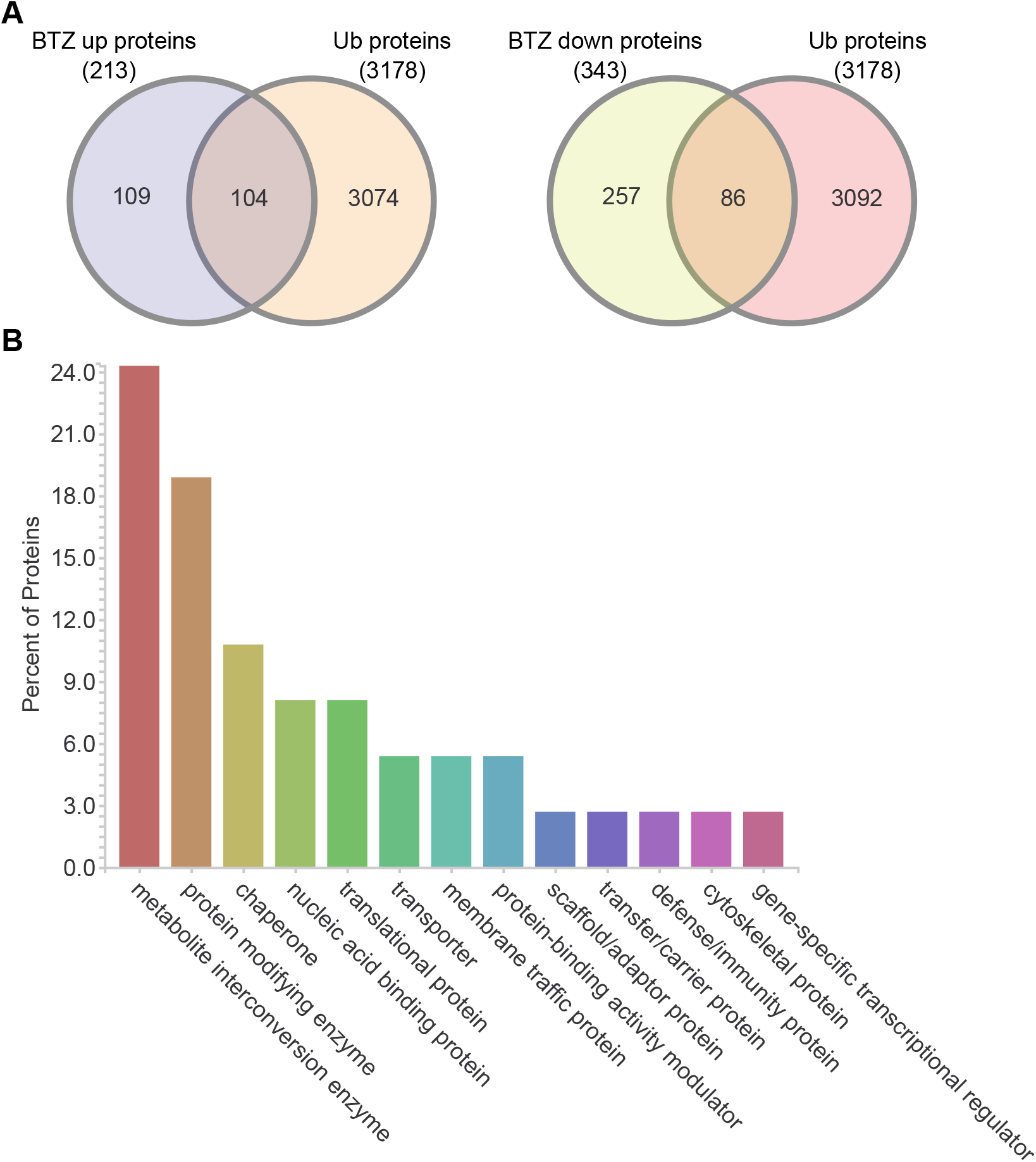
Identification of potential UPS substrates. (A) Overlap between proteins that change in abundance following BTZ treatment with proteins that are diGly modified. (B) PANTHER protein class annotations for the 104 proteins that increase in abundance following BTZ treatment and are diGly modified.

While many of these 104 proteins are potentially novel UPS substrates, several proteins which are well-established within ubiquitin mediated proteolysis pathways were observed, including CELL DIVISION CYCLE 48 (CDC48) (35), the transcription factors MYC3 (8) and AUXIN RESPONSE FACTOR 2 (ARF2) (54), the ubiquitin conjugating enzyme UBC34 (3), and the E3 ligase ABI3-INTERACTING PROTEIN 2 (AIP2) (61). These findings confirm previous studies and expand the repertoire of putative 26S proteasome substrates.

### Auxin treatment can influence ubiquitination of key auxin metabolism proteins

Ubiquitin has a longstanding and well established central role in auxin perception and signaling (19). Thus, we examined how protein abundance and diGly modified sites are impacted following 3 hours of auxin treatment (1 μM IAA). We found that 70 proteins (Supplemental Table 4) and 59 diGly modified sites (Supplemental Table 2) respond to IAA treatment (FC >1.1 p-value <0.05). Notable proteins which are ubiquitinated in response to auxin treatment include the IAA amido synthases GRETCHEN HAGAN 3.2 (GH3.2), GH3.3 (also called YDK1/BRU6/AUR3), GH3.5 (also called WES1), and GH3.6 (also called DFL1). Expression of *GH3* paralogs are classically associated with auxin response and GH3 activity directly impacts auxin homeostasis (4, 21, 45). The observed increase in GH3 protein abundance is in line with conventional wisdom that auxin induces transcription of *GH3* genes. While the auxin-induced increase in ubiquitination of GH3 proteins suggests that activation of auxin-amido conjugation enzymes via ubiquitination could occur to restore auxin homeostasis following exogenous auxin treatment.

### Motif analysis of ubiquitylated peptides identifies a novel enriched QK motif

Detailed biochemical studies of ubiquitination domains associated with protein degradation (termed “degrons”) and bioinformatic motif analyses have identified a number of conserved amino acid motifs associated with ubiquitination (12, 26, 49, 57). Using our catalog of localized diGly sites, we performed a motif analysis with motifeR to identify novel enriched motif(s) associated with ubiquitination within a 14 amino acid window. We first analyzed the diGly sites that were not increased following BTZ and found an EK^ub^ as well as several other motifs lacking any comprehensive properties (Supplemental Figure 1). A previous analysis from rice identified several enriched motifs including an EK^ub^ motif which is consistent with our results (7). We then examined the 1,336 diGly sites that are induced by BTZ and uncovered a novel QK^Ub^ motif, which is often shouldered by hydrophobic or polar amino acid residues (Figure 3A). GO enrichment analyses of QK^ub^ motif containing proteins determined that these proteins are involved in responses to plant growth regulators (auxin and karrikin), gene expression, and other biological processes (Figure 3B and Supplemental Table 5). Additionally, proteins which have the QK^Ub^ motif are significantly enriched for transcription factor molecular functions (Figure 3B).

**Fig. 3.**
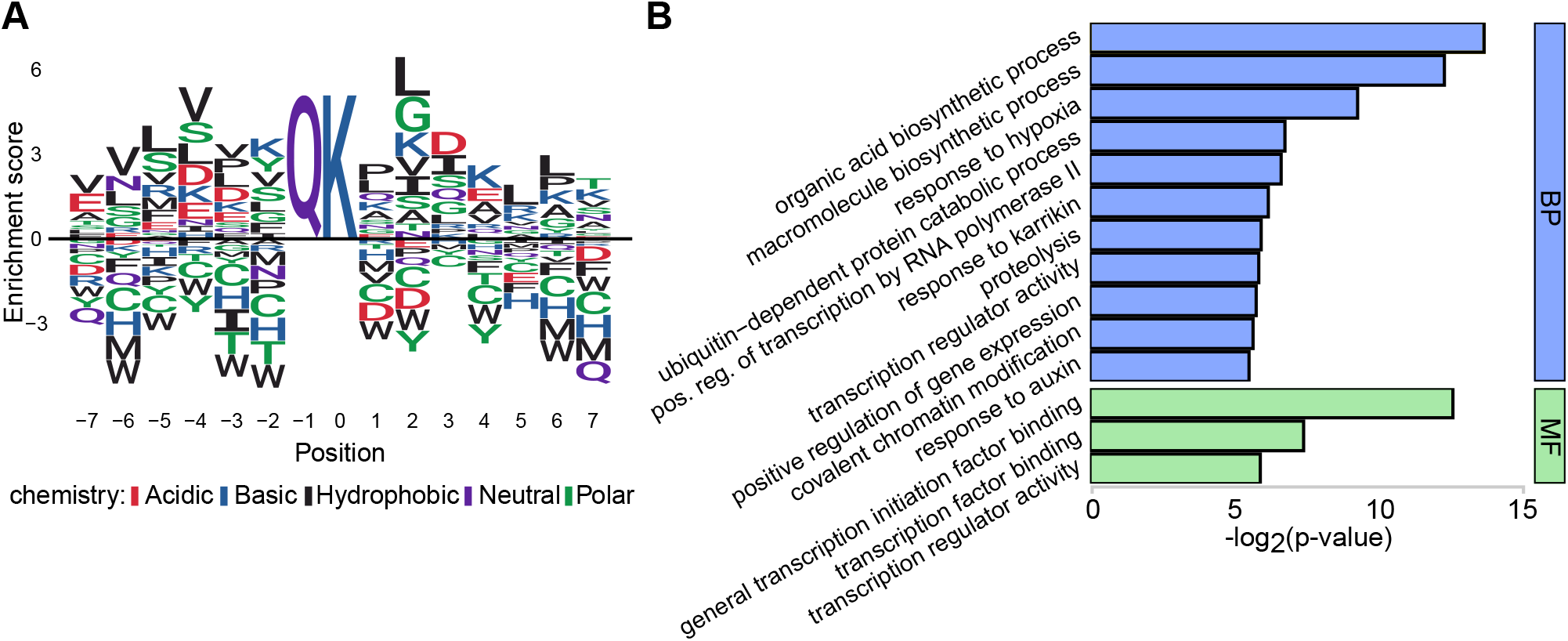
Ubiquitin motif analysis of BTZ increased diGly sites. (A) Analysis of the 14 amino acid window surrounding the 1,336 diGly sites induced by BTZ identified an enriched QK^Ub^ motif. (B) GO enrichment analyses of QK^Ub^ motif containing proteins. Supplemental Table 5 contains the full list of all identified enriched motifs surrounding ubiquitinated lysine residues.

### Ubiquitination of transcription factors is prevalent in roots and impacts protein stability

Transcription factors (TFs) were prominent among the classes of proteins detected as being ubiquitinated (Supplemental Table 2). A closer look at the modified TFs reveals that we identified over 40 different TF families with ubiquitylated proteins (Figure 4A). A number of these families contain many TFs that are modified, for example the AuxIAA, bHLH, and NAC families each contain >20 novel ubiquitination sites. Notably, many of these TFs have been well established as UPS substrates, but the site(s) of ubiquitinated have not been previously identified (23, 24). of particular interest are ubiquitination sites on twelve Aux/IAA proteins (IAA2-4, IAA7-9, IAA13, IAA16, IAA17, IAA26-28), four ARFs (ARF1, ARF2, ARF5, ARF7), six JAZ proteins (JAZ2-4, JAZ6, JAZ11, JAZ12), and three sites on ETHYLENE INSENSTIVE 3 (EIN3) (Figure 4A and Supplemental Table 2). Previous studies have identified the specific lysine residues which are ubiquitylated for paralogous proteins such as IAA6 and IAA19 (57) and sites on JAZ6 and EiN3 have been previously reported (40). These findings pinpoint *in vivo* sites of ubiquitination for many well-studied TFs involved in hormone signaling and will certainly facilitate future biochemical studies on these TFs.

**Fig. 4.**
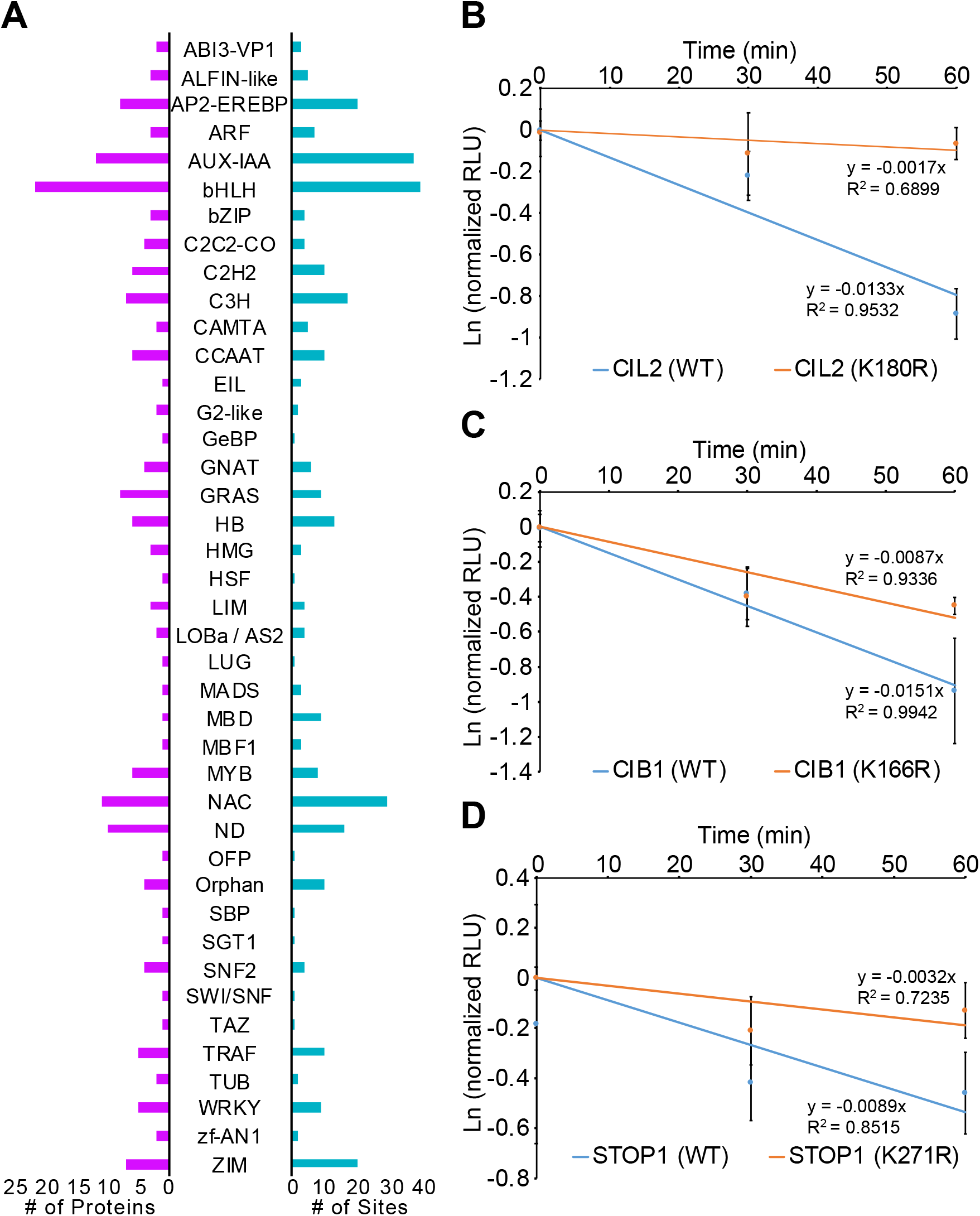
Analysis of transcription factor ubiquitination. (A) Number of ubiquitinated proteins (Left; purple) and sites (Right; blue) per transcription factor family. (B) Testing the role of UBQ attachment on protein stability *in vivo*. Degradation assays on wild-type and K>R mutated proteins. Data are means of 3 independent replicates ± standard error.

Given the key roles many transcription factors (TF) play in driving root growth, development, and environmental responses we reasoned that the observed ubiquitination of TFs may be linked to protein stability or activity. In order to test this idea we randomly selected a group of unrelated TFs for mutagenesis analyses using established luciferase based degradation assays (17). For such mutagenesis assays lysine residues are typically mutated to arginine (so called K-to-R mutants) to block ubiquitination at that particular amino acid residue (16). Using this approach WT and K>R mutated TFs were expressed as Luciferase (LUC) fusions in *Nicotiana benthamiana* and measured for protein abundance over time following cycloheximide treatment. Three of the TFs we tested displayed an increase in the K>R mutated protein compared to the WT version of the TF level in these assays (Figure 4B-D), supporting the hypothesis that these lysine residues are required for ubiquitin mediated protein turnover. This includes K166 on CRYPTOCHROME-INTERACTING BASIC-HELIX-LOOP-HELIX 1 (CIB1; AT4G34530), K180 on CIB1 LIKE PROTEIN 2 (CIL2; AT3G23690), and K271 on SENSITIVE TO PROTON RHIZOTOXICITY 1 (STOP1; AT1G34370).

Notably, all three of these TFs have expression and/or functional links to Arabidopsis root development. *STOP1* is a zinc finger TF that is required for aluminum resistance in Arabidopsis roots (13). STOP1 stability is regulated by SMALL UBIQUITIN-LIKE MODIFIER (SUMO) at K40, K212 and K395 such that blocking SUMOylation reduces stability of this TF. Additionally, *STOP1* is a component of the transcriptional regulatory network downstream of TRACHEARY ELEMENT DIFFERENTIATION iNHiBiToRY FACToR (TDIF) which is important for root vasculature development (42). *CIL2* is a bHLH transcription factor that exhibits stele enriched expression (15, 46). Like STOP1, CIL2 is also associated with the TDIF transcriptional network (42). *CIB1* is a related bHLH transcription factor that interacts with the blue light receptor CRYPTOCHROME 2 (CRY) and promotes the transition to flowering through positive regulation of FLOWERING LOCUS T (FT) (31). Based on the Interaction Viewer tool at BAR ePlant (bar.urotonto.ca/eplant/) CIB1 interacts with several other TFs with established roles in Arabidopsis root development: IAA3, IAA19, WOUND INDUCED DEDIFFERENTIATION 1 (WIND1), ULTRAPETALA (ULT), GNC-LIKE (GNL)/CYTOKININ-RPESONSIVE GATA FACTOR1 (CGA1) and HISTONE ACETYLTRANSFERASE OF THE GNAT FAMILY 1 (HAG1). The identification of ubiquitin as a key regulator of these three TFs highlights the potential regulatory role of this PTM and will support further mechanistic studies on the roles of these proteins in root growth and development.

### Final conclusions and outlook

This quantitative proteomics study provides a streamlined IP-MS/MS workflow for the identification of ubiquitination sites on plant proteins and a wealth of biochemical data for future functional follow-up assays. For example, the functional roles of the identified ubiquitination events using mutagenized proteins *in planta* can be tested using transgenic or base editing approaches. Additionally, it will be of great interest to identify the substrate-E3 ligase interactions underlying these modifications as many E3 ubiquitin ligases in Arabidopsis are still without known substrates (23, 29, 50). As future quantitative catalogs are expanded across tissues and environmental responses, we can begin to examine spatial and context-specific events that are associated with ubiquitination *in planta*.

## Materials and Methods

### Growth and treatment conditions of Arabidopsis roots

Arabidopsis Col-0 seeds were surfaced sterilized in 50 percent bleach with 0.1 percent Tween-20 for 10 minutes and rinsed five times with sterile water before plating on 0.5X Murashige and Skoog (MS) media supplemented with 1 percent sucrose and 0.8 percent agarose overlaid with pre-sterilized 110 micron nylon mesh. Col-0 seedlings were grown for 10 days in a growth chamber under 150 μmol m^-2^ s^-1^ long-day conditions at 23 C. 10 day-old Col-0 seedlings were treated with either mock solution (equal volume of DMSO solvent control), 1 μm indole-3-acetic acid (IAA; “auxin”), or 100 μm bortezomib (BTZ) at room temperature for 3 hours. Following treatments 3 g of pooled root tissue was harvested per replicate and immediately frozen into liquid nitrogen and stored at −80 C. Three independent replicates were prepared per treatment.

### Proteomic analysess

Protein extraction and digestion into peptides were done using Phenol-FASP (43, 44). Peptides containing a diGly lysine remnant were enriched using anti-diGly lysine antibody conjugated agarose beads (PTM BIO, Cat PTM-1104). Tandem Mass Tag labeling was done using TMT10plexTM label reagents (ThermoFisher, Lot TC264166) and TMTpro 16plexTM label reagents (ThermoFisher, Lot UH290430) according to a modified labeling method (44). Peptides were analyzed using an Agilent 1260 quaternary HPLC coupled to a Thermo Scientific Q-Exactive Plus high-resolution quadrupole Orbitrap mass spectrometer. All of the raw data were analyzed together using MaxQuant version 1.6.7.0. Spectra were searched against Arabidopsis TAIR10 genome, which was complemented with reverse decoy sequences and common contaminants by MaxQuant (47). For full description of proteomics methodology see supplemental methods (Supplemental File 1).

### Statistical analyses

Statistical analyses on the protein abundance and ubiquitination data were performed using TMT-NEAT Analysis Pipeline version 1.3 (https://github.com/nmclark2/TMT-Analysis-Pipeline) (10). An expanded description of statical analyses is provided in the supplemental methods (Supplemental File 1).

### Gene Ontology (GO) analyses

GO overrepresentation tests were performed using PANTHER (34) with all proteins containing a diGly site as the input and all *Arabidopsis thaliana* genes in the database as the reference (Figure 1D). The test type selected was “Fisher” with FDR correction. GO terms with a FDR < 0.05 are considered enriched. Potential UPS substrates were categorized using PANTHER protein class annotations (Figure 2B). Transcription factors were annotated obtained from (36, 59).

### Motif analyses

Motif enrichment was performed in R using the motifeR package (55). We used default settings including a 14 amino acid window size for the analysis. Lysine was set as the central residue and the TAIR10 protein annotation was used as the background reference. Sequence logo was constructed using Logolas R package and plotted using ggseqlogo R package (11, 53). Analyses were performed in R version 3.6.2 (R Core Team, 2019). GO enrichment analysis was performed using ClueGO app v2.5.7 on Cytoscape v3.8.0 (5, 41).

### Tobacco infiltration and cycloheximide treatment

Plasmids for expression WT and K>R mutated TFs are described in the supplemental methods (Supplemental File 1). Primers used for cloning are listed in Supplemental Table 6. Agrobacterium containing construct of interest was grown overnight in YEB medium supplemented with appropriate antibiotics at 28 OC. Cell pellets were resuspended and then infiltrated into *Nicotiana benthamiana* leaves. Plants were grown under light for 48 h and leaf discs were obtained from the infiltrated region of the leaves. The leaf discs were incubated in liquid MS medium containing 1 percent sucrose in the presence of 200 μM cycloheximide (CHX) or DMSO (control) for 30 and 60 minutes. The start of the first measurement following CHX treatment was designated time 0. Pooled leaf discs (approximately 60 mg) were blotted on a paper tower and flash frozen in liquid nitrogen, ground into powder and dissolved in 200 μL 1 × Cell Culture Lysis Reagent (Promega), followed by centrifugation for 15 min at 4 °C. 10 μL of the clear supernatant was used for LUC detection. The LUC activities were measured with the Luciferase Assay System (Promega) on a microplate reader. The relative light units (RLU) for each sample was analyzed as previously described (17).

## Data Availability

The original MS proteomics raw data, as well as the MaxQuant output files, may be downloaded from MassIVE (http://massive.ucsd.edu) using the identifier: MSV000086156.

## AUTHOR CONTRIBUTIONS

Conceptualization, funding acquisition, project administration, and supervision by J.W.W and D.R.K, data curation by G.S. and N.M.C, formal analysis by D.O., N.M.C., and C.M., investigation and validation by G.S., D.O. and Y.P., methodology by G.S., D.O., C.M., Y.P, D.R.K., and J.W.W., software by N.M.C., visualization and writing by J.W.W., D.R.K., D.O., and C.M.

## ACKNOWLEDGEMENTS

This work was supported by the Iowa State University Plant Science Institute, NIH (R01GM120316), NSF (IOS-1759023) and USDA NIFA Hatch project IOW3808 funds to J.W.W.; USDA NIFA Hatch project IOW3649 and ISU start-up funds to D.R.K. N.M.C. is supported by a USDA NIFA Postdoctoral Research Fellowship (2019-67012-29712).

This manuscript was formatted in Overleaf using the Henriques Lab bioRxiv template.

## References

1. Abu Hatoum, O., Gross-Mesilaty, S., Breitschopf, K., Hoffman, A., Gonen, H., Ciechanover, A., and Bengal, E. Degradation of myogenic transcription factor MyoD by the ubiquitin pathway in vivo and in vitro: Regulation by specific DNA binding. Molecular and Cellular Biology 18, 10, 5670–5677.

2. Aguilar-Hernández, V., Kim, D.-Y., Stankey, R.J., Scalf, M., Smith, L.M., and Vierstra, R.D. Mass spectrometric analyses reveal a central role for ubiquitylation in remodeling the arabidopsis proteome during photomorphogenesis. Molecular Plant 10, 6, 846–865.

3. Ahn, M.Y., Oh, T.R., Seo, D.H., Kim, J.H., Cho, N.H., and Kim, W.T. Arabidopsis group XIV ubiquitin-conjugating enzymes AtuBC32, AtuBC33, and AtuBC34 play negative roles in drought stress response. Journal of Plant Physiology 230, 73–79.

4. Aoi, Y., Tanaka, K., Cook, S.D., Hayashi, K.-I., and Kasahara, H. GH3 auxin-amido synthetases alter the ratio of indole-3-acetic acid and phenylacetic acid in arabidopsis. Plant and Cell Physiology 61, 3, 596–605.

5. Bindea, G., Mlecnik, B., Hackl, H., Charoentong, P., Tosolini, M., Kirilovsky, A., Fridman, W.-H., Pagès, F., Trajanoski, Z., and Galon, J. ClueGO: a cytoscape plug-in to decipher functionally grouped gene ontology and pathway annotation networks. Bioinformatics (Oxford, England) 25, 8, 1091–1093.

6. Cadwell, K. Ubiquitination on nonlysine residues by a viral e3 ubiquitin ligase. Science 309, 5731, 127–130.

7. Chen, X.-L., Xie, X., Wu, L., Liu, C., Zeng, L., Zhou, X., Luo, F., Wang, G.-L., and Liu, W. Proteomic analysis of ubiquitinated proteins in rice (oryza sativa) after treatment with pathogen-associated molecular pattern (PAMP) elicitors. Frontiers in Plant Science 9. Publisher: Frontiers.

8. Chico, J.M., Lechner, E., Fernandez-Barbero, G.,Canibano, E., García-Casado, G., Franco-Zorrilla, J.M., Hammann, P., Zamarreño, A.M., García-Mina, J.M., Rubio, V., Genschik, P., and Solano, R. CUL3bpm e3 ubiquitin ligases regulate MYC2, MYC3, and MYC4 stability and JA responses. Proceedings of the National Academy of Sciences 117,11,6205–6215.

9. Ciechanover, A. N-terminal ubiquitination: more protein substrates join in. Trends in Cell Biology 14,3, 103–106.

10. Clark, N.M., Nolan, T.M., Wang, P., Song, G., Montes, C., Guo, H., Sozzani, R., Yin, Y., and Walley, J.W. Integrated omics networks reveal the temporal signaling events of brassinosteroid response in Arabidopsis. bioRxiv.

11. Dey, K.K., Xie, D., and Stephens, M. A new sequence logo plot to highlight enrichment and depletion. BMC Bioinformatics 19, 1, 473.

12. Dreher, K.A., Brown, J., Saw, R.E., and Callis, J. The arabidopsis aux/IAA protein family has diversified in degradation and auxin responsiveness. The Plant Cell 18, 3, 699–714.

13. Fang, Q., Zhang, J., Zhang, Y., Fan, N., van den Burg, H.A., and Huang, C.-F. Regulation of aluminum resistance in arabidopsis involves the SUMOylation of the zinc finger transcription factor STOP1. The Plant Cell 32, 12, 3921–3938.

14. Fulzele, A., and Bennett, E.J. Ubiquitin diGLY proteomics as an approach to identify and quantify the ubiquitin-modified proteome. in The Ubiquitin Proteasome System, T. Mayor and G. Kleiger, Eds., vol. 1844. Springer New York, pp. 363–384. Series Title: Methods in Molecular Biology.

15. Gaudinier, A., Zhang, L., Reece-Hoyes, J.S., Taylor-Teeples, M., Pu, L., Liu, Z., Breton, G., Pruneda-Paz, J.L., Kim, D., Kay, S.A., Walhout, A.J.M., Ware, D., and Brady,S. M. Enhanced y1h assays for arabidopsis. Nature Methods 8,12, 1053–1055.

16. Gilkerson, J., Kelley, D.R., Tam, R., Estelle, M., and Callis, J. Lysine residues are not required for proteasome-mediated proteolysis of the auxin/indole acidic acid protein IAA1. Plant Physiology 168, 2, 708–720.

17. Gilkerson, J., Tam, R., Zhang, A., Dreher, K., and Callis,J. Cycloheximide assays to measure protein degradation in vivo in plants. BIO-PROTOCOL 6, 17.

18. Gladman, N.P., Marshall, R.S., Lee, K.-H., and Vierstra, R.D. The proteasome stress regulon is controlled by a pair of NAC transcription factors in arabidopsis. The Plant Cell 28, 6, 1279–1296.

19. Gray, W.M., and Estelle, M. Function of the ubiquitin–proteasome pathway in auxin response. Trends in Biochemical Sciences 25, 3, 133–138.

20. Grubb, L.E., Derbyshire, P., Dunning, K., Zipfel, C., Menke,F. L. H., and Monaghan, J. Large-scale identification of ubiquitination sites on membrane-associated proteins in arabidopsis thaliana seedlings. bioRxiv, 2020.09.16.299883. Publisher: Cold Spring Harbor Laboratory Section: New Results.

21. Hagen, G., and Guilfoyle, T.J. Rapid induction of selective transcription by auxins. Molecular and Cellular Biology 5, 6, 1197–1203.

22. Igawa, T., Fujiwara, M., Takahashi, H., Sawasaki, T., Endo,Y., Seki, M., Shinozaki, K., Fukao, Y., and Yanagawa, Y. Isolation and identification of ubiquitin-related proteins from arabidopsis seedlings. Journal of Experimental Botany 60, 11, 3067–3073. Publisher: oxford Academic.

23. Kelley, D.R. E3 ubiquitin ligases: Key regulators of hormone signaling in plants. Molecular& Cellular Proteomics 17, 6, 1047–1054.

24. Kelley, D.R., and Estelle, M. Ubiquitin-mediated control of plant hormone signaling. Plant Physiology 160, 1,47–55.

25. Kim,D.-Y., Scalf, M., Smith, L.M., and Vierstra, R.D. Advanced proteomic analyses yield a deep catalog of ubiquitylation targets in arabidopsis. The Plant Cell 25, 5, 1523–1540. Publisher: American Society of Plant Biologists Section: LARGE-SCALE BIOLOGY articles.

26. Kim, W., Bennett, E.J., Huttlin, E.L., Guo, A., Li, J., Possemato, A., Sowa, M.E., Rad, R., Rush, J., Comb, M.J., Harper, J.W., and Gygi, S.P. Systematic and quantitative assessment of the ubiquitin-modified proteome. Molecular cell 44, 2, 325–40.

27. Kraft, E., Stone, S.L., Ma, L., Su, N., Gao, Y., Lau, O.-S., Deng, X.-W., and Callis, J. Genome analysis and functional characterization of the e2 and RING-type e3 ligase ubiquitination enzymes of arabidopsis. Plant Physiology 139,4, 1597–1611.

28. Kravtsova-Ivantsiv, Y., and Ciechanover, A. Non-canonical ubiquitin-based signals for proteasomal degradation. Journal of Cell Science 125, 3, 539–548.

29. Lee, C.-M., Feke, A., Li, M.-W., Adamchek, C., Webb, K., Pruneda-Paz, J., Bennett, E.J., Kay, S.A., and Gendron,J. M. Decoys untangle complicated redundancy and reveal targets of circadian clock f-box proteins. Plant Physiology 177, 3, 1170–1186.

30. Li, X.-M., Chao, D.-Y., Wu, Y., Huang, X., Chen, K., Cui, L.-G., Su, L., Ye, W.-W., Chen, H., Chen, H.-C., Dong, N.-Q., Guo,T., Shi, M., Feng, Q., Zhang, P., Han, B., Shan, J.-X., Gao, J.-P., and Lin, H.-X. Natural alleles of a proteasome α2 subunit gene contribute to thermotolerance and adaptation of african rice. Nature Genetics 47, 7, 827–833. Number: 7 Publisher: Nature Publishing Group.

31. Liu, Y., Li, X., Ma, D., Chen, Z., Wang, J., and Liu, H. CIB1 and CO interact to mediate CRY2-dependent regulation of flowering. EMBO reports 19, 10.

32. Manzano, C., Abraham, Z., López-Torrejón, G., and Del Pozo, J.C. Identification of ubiquitinated proteins in arabidopsis. Plant Molecular Biology 68, 1, 145–158.

33. Maor, R., Jones, A., Nühse, T.S., Studholme, D.J., Peck, S.C., and Shirasu, K. Multidimensional protein identification technology (MudPiT) analysis of ubiquitinated proteins in plants. Molecular & Cellular Proteomics 6, 4, 601–610. Publisher: American Society for Biochemistry and Molecular Biology Section: Research.

34. Mi, H.,Muruganujan, A., Ebert, D., Huang, X., and Thomas,P. D. PANTHER version 14: more genomes, anew PANTHER GO-slim and improvements in enrichment analysis tools. Nucleic Acids Research 47, D419–D426. Publisher: Oxford Academic.

35. Mérai, Z., Chumak, N., García-Aguilar, M., Hsieh, T.-F., Nishimura, T., Schoft, V.K., Bindics, J., Slusarz, L., Arnoux, S., Opravil, S., Mechtler, K., Zilberman, D., Fischer, R.L., and Tamaru, H. The AAA-ATPase molecular chaperone cdc48/p97 disassembles sumoylated centromeres, decondenses heterochromatin, and activates ribosomal RNA genes. Proceedings of the National Academy of Sciences 111, 45, 16166–16171.

36. Pruneda-Paz, J.L., Breton, G., Nagel, D.H., Kang, S.E., Bonaldi, K., Doherty, C.J., Ravelo, S., Galli, M., Ecker,J. R., and Kay, S.A. A genome-scale resource for the functional characterization of arabidopsis transcription factors. Cell Reports 8,2, 622–632.

37. Romero-Barrios, N., and Vert, G. Proteasome-mindependent functions of lysine-63 polyubiquitination in plants. New Phytologist 217,3, 995–1011. _eprint: https://nph.onlinelibrary.wiley.com/doi/pdf/10.1111/nph.14915.

38. Rose, C.M., Isasa, M., Ordureau, A., Prado, M.A., Beausoleil, S.A., Jedrychowski, M.P., Finley, D.J., Harper,J. W., and Gygi, S.P. Highly multiplexed quantitative mass spectrometry analysis of ubiquitylomes. Cell Systems 3, 4, 395–403.e4. Publisher: Elsevier.

39. Santner, A., and Estelle, M. The ubiquitin-proteasome system regulates plant hormone signaling. The Plant Journal 61, 6, 1029–1040.

40. Saracco, S.A., Hansson, M., Scalf, M., Walker, J.M., Smith, L.M., and Vierstra, R.D. Tandem affinity purification and mass spectrometric analysis of ubiquitylated proteins in arabidopsis. The Plant Journal: For Cell and Molecular Biology 59, 2, 344–358.

41. Shannon, P., Markiel, A., Ozier, O., Baliga, N.S., Wang,J. T., Ramage, D., Amin, N., Schwikowski, B., and Ideker,T. cytoscape: a software environment for integrated models of biomolecular interaction networks. Genome Res 13, 11, 2498–2504.

42. Smit, M.E., McGregor, S.R., Sun, H., Gough, C., Bågman,A.-M., Soyars, C.L., Kroon, J.T., Gaudinier, A., Williams,C. J., Yang, X., Nimchuk, Z.L., Weijers, D., Turner, S.R., Brady, S.M., and Etchells, J.P. A PXY-mediated transcriptional network integrates signaling mechanisms to control vascular development in arabidopsis. The Plant Cell 32, 2, 319–335.

43. Song, G., Hsu, P.Y., and Walley, J.W. Assessment and refinement of sample preparation methods for deep and quantitative plant proteome profiling. PROTEOMICS 18, 17, 1800220. Publisher: Wiley-Blackwell.

44. Song, G., Montes, C., and Walley, J.W. Quantitative profiling of protein abundance and phosphorylation state in plant tissues using tandem mass tags. in Plant Proteomics: Methods and Protocols, J. V. Jorrin-Novo, L. Valledor, M.A. Castillejo, and M.-D. Rey, Eds., Methods in Molecular Biology. Springer US, pp. 147–156.

45. Staswick, P.E., Serban, B., Rowe, M., Tiryaki, I., Maldonado, M.T., Maldonado, M.C., and Suza, W. Characterization of an arabidopsis enzyme family that conjugates amino acids to indole-3-acetic acid. The Plant Cell 17, 2, 616–627.

46. Taylor-Teeples, M., Lin, L., de Lucas, M., Turco, G., Toal,T. W., Gaudinier, A., Young, N.F., Trabucco, G.M., Veling,M. T., Lamothe, R., Handakumbura, P.P., Xiong, G., Wang,C., Corwin, J., Tsoukalas, A., Zhang, L., Ware, D., Pauly,M., Kliebenstein, D.J., Dehesh, K., Tagkopoulos, I., Breton, G., Pruneda-Paz, J.L., Ahnert, S.E., Kay, S.A., Hazen,S. P., and Brady, S.M. An arabidopsis gene regulatory network for secondary cell wall synthesis. Nature 517, 7536, 571–575.

47. Tyanova, S., Temu, T., and Cox, J. The MaxQuant computational platform for mass spectrometry-based shotgun proteomics. Nature Protocols 11, 12, 2301–2319. Publisher: Nature Research.

48. Udeshi, N.D., Mani, D.C., Satpathy, S., Fereshetian, S., Gasser, J.A., Svinkina, T., Olive, M.E., Ebert, B.L., Mertins, P., and Carr, S.A. Rapid and deep-scale ubiquitylation profiling for biology and translational research. Nature Communications 11, 1, 1–11. Number: 1 Publisher: Nature Publishing Group.

49. Varshavsky, A. Naming atargeting signal. Cell 64, 1, 13–15.

50. Vierstra, R.D. The expanding universe of ubiquitin and ubiquitin-like modifiers. Plant Physiology 160, 1,2–14.

51. Vierstra, R.D. The ubiquitin-26s proteasome system at the nexus of plant biology. Nature Reviews Molecular Cell Biology 10,6, 385–397. Number: 6 Publisher: Nature Publishing Group.

52. Vissenberg, K., Claeijs, N., Balcerowicz, D., and Schoenaers, S. Hormonal regulation of root hair growth and responses to the environment in arabidopsis. Journal of Experimental Botany 71, 8, 2412–2427.

53. Wagih, O. ggseqlogo: a versatile r package for drawing sequence logos. Bioinformatics (Oxford, England) 33, 22, 3645–3647.

54. Walton, A., Stes, E., Cybulski, N., Bel, M.V., Iñigo, S., Durand, A.N., Timmerman, E., Heyman, J., Pauwels, L., Veylder, L.D., Goossens, A., Smet, I.D., Coppens, F., Goormachtig, S., and Gevaert, K. It’s time for some “site”-seeing: Novel tools to monitor the ubiquitin landscape in arabidopsis thaliana. The Plant Cell 28, 1,6–16. Publisher: American Society of Plant Biologists Section: Perspective.

55. Wang, S., Cai, Y., Cheng, J., Li, W., Liu, Y., and Yang, H. motifeR: An integrated web software for identification and visualization of protein posttranslational modification motifs. Proteomics 19, 23, e1900245.

56. Wang, Y.-F., Chao, Q., Li, Z., Lu, T.-C., Zheng, H.-Y., Zhao, C.-F., Shen, Z., Li, X.-H., and Wang, B.-C. Large-scale identification and time-course quantification of ubiquitylation events during maize seedling de-etiolation. Genomics, Proteomics& Bioinformatics 17, 6, 603–622.

57. Winkler, M., Niemeyer, M., Hellmuth, A., Janitza, P., Christ, G., Samodelov, S.L., Wilde, V., Majovsky, P., Trujillo, M., Zurbriggen, M.D., Hoehenwarter, W., Quint, M., and Calderón Villalobos, L.I. A. Variation in auxin sensing guides AUX/IAA transcriptional repressor ubiquitylation and destruction. Nature Communications 8, 1, 15706.

58. Xie, X., Kang, H., Liu, W., and Wang, G.-L. Comprehensive profiling of the rice ubiquitome reveals the significance of lysine ubiquitination in young leaves. Journal of Proteome Research 14, 5, 2017–2025. Publisher:American Chemical Society

59. Yilmaz, A., Nishiyama, M.Y., Fuentes, B.G., Souza, G.M., Janies, D., Gray, J., and Grotewold, E. GRASSIUS: A platform for comparative regulatory genomics across the grasses. Plant Physiol. 149, 1, 171–180.

60. Zhang, N., Zhang, L., Shi, C., Tian, Q., Lv, G., Wang, Y., Cui, D., and Chen, F. Comprehensive profiling of lysine ubiquitome reveals diverse functions of lysine ubiquitination in common wheat. Scientific Reports 7, 1, 13601. Number: 1 Publisher: Nature Publishing Group.

61. Zhang, X. The AIP2 e3 ligase acts as a novel negative regulator of ABA signaling by promoting ABi3 degradation. Genes& Development 19, 13, 1532–1543.

