## Supplemental File 1 for "Quantitative proteomics reveals extensive lysine ubiquitination in the Arabidopsis root proteome and uncovers novel transcription factor stability states"

**Supplemental Materials and Methods**

### Protein extraction and proteomics analyses

Protein extraction and digestion were done using Phenol-FASP methods (Song et al., 2018; Song et al., 2020). For this, Arabidopsis root tissue was ground under liquid nitrogen for 15 minutes. Next, 5 volumes of Tris buffered phenol pH 8 (buffer:tissue, v:w) was added to each sample tube which was vortexed for 1 minute. The samples were then vortexed for 1 more minute with 5 volumes (buffer:tissue, v:w) of extraction buffer (50 mM Tris pH 7.5, 1 mM EDTA pH 8, 0.9 M sucrose, 1x phosphatase inhibitors, 10mM N-Ethylmaleimide (NEM, Cat# 156100100) ), and then centrifuged at 13,000 x g for 10 min at 4°C. The phenol phase was transferred to a new tube and a second phenol extraction was performed on the aqueous phase. The two phenol phase extractions were combined and 5 volumes of prechilled methanol with 0.1 M ammonium acetate was added and mixed well, then kept at -80°C for 1h prior to centrifugation at 4,500 x g for 10 min at 4°C. Precipitation with 0.1 M ammonium acetate in methanol was performed twice with incubation at -20°C for 30 min. The sample was resuspended in 70% methanol and kept at -20°C for 30 min prior to centrifuging at 4,500 x g, for 10 min at 4°C. The supernatant was discarded, and the pellet was placed in a vacuum concentrator till near dry. Two volumes (buffer:pellet, v:v) of protein extraction buffer (8M urea, 50 mM Tris pH 7, 5 mM TCEP, 1x phosphatase inhibitors, 10mM NEM) was added to the pellet. The samples were then probe sonicated to aid in resuspension of the pellet. The protein concentration was then determined using the Bradford assay (Thermo Scientific).

*Filter Aided Sample Preparation (FASP)*

The solubilized protein (~2.5 mg in 0.5 ml) was mixed with 3.5 ml urea solution (8 M urea, 100 mM Tris pH 8, 1x phosphatase inhibitors, 10 mM NEM), added to an Amicon Ultracel – 30K centrifugal filter (Cat # UFC803008), and centrifuged at 4,000 x g for 40-60 min. This step was repeated once. Then, 4 ml of urea solution with 2 mM TCEP was added to the filter unit and centrifuged at 4,000 x g for 20-40 min. Next, 2 ml IAM solution (50 mM iodoacetamide in urea solution) was added and incubated without mixing at room temperature for 30 min in the dark prior to centrifuging at 4,000 x g for 20-40 min. Two ml of urea solution was added to the filter unit, which was then centrifuged at 4,000 x g for 20-40 min. This step was repeated once. Two ml of 0.05 M $NH_4HCO_3$ with 1x phosphatase inhibitors and 10 mM NEM was added to the filter unit and centrifuged at 4,000 x g for 20-40 min. This step was repeated once. Then 2 ml of 0.05 M $NH_4HCO_3$ solution with trypsin (enzyme to protein ratio 1:100) was added. Samples were incubated at 37°C overnight. Undigested protein was estimated using Bradford assays, then trypsin (1µg/µl) was added to a ratio of 1:100 and an equal volume of Lys-C (0.1 µg/µl) were added to the sample and incubated for an additional 4 hours at 37°C. The filter unit was added to a new collection tube and centrifuged at 4,000 x g for 20-40 min. One ml of 0.05M $NH_4HCO_3$ was added and centrifuged at 4,000 x g for 20-40 min. The samples were acidified to pH 2-3 with 100% formic acid and centrifuged at 21,000 x g for 20 min. Finally, samples were desalted using 100 mg or 500 mg Sep-Pak C18 cartridges (Waters). Eluted peptides were dried using a vacuum centrifuge (Thermo) and resuspended in 0.1% formic acid. Peptide amount was quantified using the Pierce BCA Protein assay kit.

*Di-glycine lysine peptide enrichment*

Peptides containing a diGly lysine remnant were enriched using anti-diGly lysine antibody conjugated agarose beads (PTM BIO, Cat# PTM-1104). In each experiment below, anti-diGly lysine antibody conjugated agarose beads were prewashed 3 times using 0.5 ml ice-cold phosphate buffered saline (PBS).

For the small-scale 1D-MS/MS pilot experiment, 2 mg of C18 desalted peptides resuspended in 400 $\mu$l IP/Wash buffer were combined with 40 $\mu$l of PBS pre-washed anti-diGly lysine antibody-conjugated agarose beads and rotated overnight at 4°C. The next day, the sample was centrifuged at 500 g for 1 minute and the supernatant was removed. The beads were then washed 3 times by adding 0.5 ml IP/Wash buffer (100mM NaCl, 1mM EDTA, 20mM Tris-HCl, PH8), mixed by inverting for 30 seconds, and then centrifuging at 500 g for 1 minute. After the last wash step, 0.5 ml of 0.1% trifluoroacetic acid (TFA) was added, which was followed by 1 minute gentle rotation at room temperature to elute the bound peptides. The eluted peptides were collected by centrifugation at 1,000 g for 1 minute at room temperature, then transferred to a new tube. The elution steps were repeated two more times. Note that for the last elution step, the beads were rotated in 0.1% TFA for 5 minutes instead 1 minute. The eluted peptides from each step were combined together, vacuum centrifuged to dry, and stored at -80ºC

For the "2D-label free" experiment (Figure 1A&C), 20 mg C18 desalted peptides and 400 $\mu$l anti-diGly lysine antibody conjugated agarose beads were used. Specifically, the 20 mg peptides were resuspended with 3.6 ml IP/Wash buffer and then 600 $\mu$l of the resuspended peptides were aliquoted to individual 1.5 ml tubes. Next, 66.6 $\mu$l of PBS pre-washed anti-diGly lysine antibody conjugated agarose beads were added to each tube and rotated overnight at 4°C. The beads were washed and peptides eluted as described above. Eluted peptides were stored at -80ºC.

For the "Enrich -> TMT" experiments (Figure 1A&C), 2 mg of C18 desalted peptides of each sample (9 samples total) were resuspended with 400 $\mu$l IP/Wash buffer, combined with 40 $\mu$l of PBS pre-washed anti-diGly lysine antibody conjugated agarose beads and, rotated overnight at 4°C. The beads were washed and peptides eluted as described above. Eluted peptides were stored at -80ºC until TMT labeling.

For the "TMT -> Enrich" test (Figure 1A), 1.8 mg of desalted peptides, per sample (9 samples total), were labeled with TMT10plex labeling reagents. Each 1.8 mg sample was labeled with 2.4 mg TMT reagent as described below. The TMT labeled peptides were pooled. The 18 mg of pooled TMT labeled peptides were desalted using 500 mg Sep-Pak C18 cartridges (Waters) and enriched using 300 $\mu$l of anti-diGly lysine antibody as described above.

**TMT Labeling**

TMT10plex™ label reagents (ThermoFisher, Lot #TC264166) and TMTpro 16plex™ label reagents (ThermoFisher, Lot #UH290430) were used to label the anti di-Gly lysine antibody enriched peptides according to a modified labeling method (Song et al., 2020). The diGly enriched samples were labeled as follows. All peptides that were recovered following diGly enrichment, from each sample, were resuspended with 20 $\mu$l of 0.2 M HEPES buffer pH 8.5 (Alfa Aesar Cat# J63218) and then mixed with 0.08 mg TMT or TMTpro reagent that was resuspended in 8 $\mu$l dry acetonitrile (Millipore cat# AX0143-7). After 2-hour incubation at room temperature, 1.6 $\mu$l of 5% hydroxylamine were added to each tube and vortexed. The samples were incubated at room temperature for 15 minutes to quench the labeling reaction. Next, the 9 samples were mixed together and stored at -80ºC.

For the TMTpro protein abundance runs, 10 $\mu$g of C18 desalted peptides were resuspended in 20 $\mu$l of 0.2 M HEPES buffer pH 8.5 and then mixed with 0.08mg TMT reagent that was resuspended in 8 $\mu$l dry acetonitrile. After 2-hour incubation at room temperature, 1.6 $\mu$l of 5% hydroxylamine were added to each tube and vortexed. The samples were incubated at room temperature for 15 minutes to quench the labeling reaction. Next, the 9 samples were mixed together and stored at -80ºC.

## LC/MS-MS

An Agilent 1260 quaternary HPLC was used to deliver a flow rate of ~300 or 600 nL per minute for 100 or 200 ID packed nanospray emitter tips respectively, via a splitter. All columns were packed in house using a Next Advance pressure cell and the nanospray tips were fabricated using fused silica capillary that was pulled to a sharp tip using a laser puller (Sutter P-2000).

*Small-scale 1D-MS/MS pilot experiment*

All of the diGly lysine enriched peptides (from 2 mg input peptides) were loaded onto 5 cm capillary columns packed with 5 μM Zorbax SB-C18 (Agilent), which was connected using a zero dead volume 1 μm filter (Upchurch, M548) to a 20 cm nanospray tip (200 μm ID) packed with 2.5 μM C18 (Waters). Peptides were separated and delivered to the mass spectrometer using a 150 min reverse-phase gradient comprised of 5-30% (ACN, 0.1% formic acid) over 120 min, 30-80% (ACN, 0.1% formic acid) over 20 min, 80% (ACN, 0.1% formic acid) 5 min hold, and 80-0% (ACN, 0.1% formic acid) over 5 min.

Eluted peptides were analyzed using a Thermo Scientific Q-Exactive Plus high-resolution quadrupole Orbitrap mass spectrometer, which was directly coupled to the HPLC. Data dependent acquisition was obtained using Xcalibur 4.0 software in positive ion mode with a spray voltage of 2.00 kV and a capillary temperature of 275 °C and an RF of 60. MS1 spectra were measured at a resolution of 70,000, an automatic gain control (AGC) of 3e6 with a maximum ion time of 100 ms and a mass range of 400-2000 m/z. Up to 15 MS2 were triggered at a resolution of 17,500 with a fixed first mass of 120 m/z. An AGC of 1e5 with a maximum ion time of 50 ms, an isolation window of 1.3 m/z, and a normalized collision energy of 28 were used for this run. Charge exclusion was set to unassigned, 1, 5–8, and >8. MS1 that triggered MS2 scans were dynamically excluded for 25 s.

*2D-label free LC-MS/MS*

All of the di-Gly enriched peptides were loaded onto 10 cm capillary columns packed with 5 μM Zorbax SB-C18 (Agilent), which was connected using a zero dead volume 1 μm filter (Upchurch, M548) to a 5 cm long strong cation exchange (SCX) column packed with 5 μm PolySulfoethyl (PolyLC). The SCX column was then connected to a 20 cm long nanospray tip (100 μm ID) packed with 2.5 μM C18 (Waters). The 3 sections were joined and mounted on a custom electrospray source for on-line nested peptide elution. Peptides were eluted from the loading column unto the SCX column using a 0 to 80% acetonitrile gradient over 60 minutes. Peptides were then fractionated from the SCX column using a series of salt steps. The following ammonium acetate salt steps were used: 25, 45, 60, 70, 80, 90, 100, 300, 500 and 1000 mM. For these analyses, buffers A (99.9% $H_2O$, 0.1% formic acid), B (99.9% ACN, 0.1% formic acid), C (100 mM ammonium acetate, 2% formic acid), and D (1 M ammonium acetate, 2% formic acid) were utilized. For each salt step, a 150-minute gradient program comprised of a 0–5 minute increase to the specified ammonium acetate concentration, 5–10 minutes hold, 10–14 minutes at 100% buffer A, 15–100 minutes 5–30% buffer B, 100-121 minutes 30–45% buffer B, 120–140 minutes 45–80% buffer B, 140–144 minutes 80% buffer B, and 145–150 minutes buffer A was employed.

Eluted peptides were analyzed using a Thermo Scientific Q-Exactive Plus high-resolution quadrupole Orbitrap mass spectrometer using acquisition setting as described above for the 1D-MS/MS analysis.

*"Enrich -> TMT" 2D-TMT LC-MS/MS*

All of the TMT labeled diGly lysine enriched peptides were loaded onto 10 cm capillary columns packed with 5 μM Zorbax SB-C18 (Agilent), which was connected using a zero dead volume 1 μm filter (Upchurch, M548) to a 5 cm long strong cation exchange (SCX) column packed with 5 μm PolySulfoethyl (PolyLC). The SCX column was then connected to a 20 cm nanospray tip (200 μm ID) packed with 2.5 μM C18 (Waters). The 3 sections were joined and mounted on a custom electrospray source for on-line nested peptide elution. Peptides were eluted from the loading column unto the SCX column using a 0 to 80% acetonitrile gradient over 60 minutes. Peptides were then fractionated from the SCX column using these ammonium acetate salt steps: 35, 70, 100 and 1000 mM.

Eluted peptides were analyzed using a Thermo Scientific Q-Exactive Plus high-resolution quadrupole Orbitrap mass spectrometer, which was directly coupled to the HPLC. Data dependent acquisition was obtained using Xcalibur 4.0 software in positive ion mode with a spray voltage of 2.00 kV and a capillary temperature of 275 °C and an RF of 60. MS1 spectra were measured at a resolution of 70,000, an automatic gain control (AGC) of 3e6 with a maximum ion time of 100 ms and a mass range of 400-2000 m/z. Up to 15 MS2 were triggered at a resolution of 35,000 with a fixed first mass of 120 m/z. An AGC of 1e5 with a maximum ion time of 50 ms, an isolation window of 1.3 m/z, and a normalized collision energy of 33 were used for this analysis. Charge exclusion was set to unassigned, 1, 5–8, and >8. MS1 that triggered MS2 scans were dynamically excluded for 25 s.

*"Enrich -> TMTpro" 2D-TMTpro LC-MS/MS*

All of the TMTpro labeled diglycine lysine enriched peptides were loaded onto 10 cm capillary columns packed with 5 μM Zorbax SB-C18 (Agilent), which was connected using a zero dead volume 1 μm filter (Upchurch, M548) to a 5 cm long strong cation exchange (SCX) column packed with 5 μm PolySulfoethyl (PolyLC). The SCX column was then connected to a 20 cm nanospray tip (100 μm ID) packed with 2.5 μM C18 (Waters). The 3 sections were joined and mounted on a custom electrospray source for on-line nested peptide elution. Peptides were eluted from the loading column unto the SCX column using a 0 to 80% acetonitrile gradient over 60 minutes. Peptides were then fractionated from the SCX column using a series of salt steps. The following ammonium acetate salt steps were used: 30, 60, 80, 90, 95, 100, 200 and 1000 mM.
Eluted peptides were analyzed using a Thermo Scientific Q-Exactive Plus high-resolution quadrupole Orbitrap mass spectrometer using acquisition settings described in *"Enrich -> TMT" 2D-TMT LC-MS/MS"* except a normalized collision energy of 31 were used for this analysis.

*"TMTpro protein abundance analysis"*

Ten $\mu$g of TMTpro labeled peptides were loaded onto 10 cm capillary columns packed with 5 μM Zorbax SB-C18 (Agilent), which was connected using a zero dead volume 1 μm filter (Upchurch, M548) to a 5 cm long strong cation exchange (SCX) column packed with 5 μm PolySulfoethyl (PolyLC). The SCX column was then connected to a 20 cm nanospray tip (100 μm ID) packed with 2.5 µM C18 (Waters). The 3 sections were joined and mounted on a custom electrospray source for on-line nested peptide elution. Peptides were eluted from the loading column unto the SCX column using a 0 to 80% acetonitrile gradient over 60 minutes. Peptides were then fractionated from the SCX column using a series of salt steps. The following ammonium acetate salt steps were used: 30, 60, 70, 75, 80, 82.5, 85, 87.5, 90, 92.5, 95, 97.5, 100, 125, 150, 200 and 1000 mM.

Eluted peptides were analyzed using a Thermo Scientific Q-Exactive Plus high-resolution quadrupole Orbitrap mass spectrometer, which was directly coupled to the HPLC. Data dependent acquisition was obtained using Xcalibur 4.0 software in positive ion mode with a spray voltage of 2.2 kV and a capillary temperature of 275 °C and an RF of 60. MS1 spectra were measured at a resolution of 70,000, an automatic gain control (AGC) of 3e6 with a maximum ion time of 100 ms and a mass range of 400-2000 m/z. Up to 15 MS2 were triggered at a resolution of 17,500 or 35,000 was used for two replicate runs respectively. Note that the 9 TMTpro labels used here have 1 Da spacing between reporter ion, which enables acquisition with the 17,500 resolution setting. A fixed first mass of 120 m/z. An AGC of 1e5 with a maximum ion time of 50 ms, an isolation window of 1.3 m/z, and a normalized collision energy of 31 were used. Charge exclusion was set to unassigned, 1, 5–8, and >8. MS1 that triggered MS2 scans were dynamically excluded for 25 s.

*Database search and FDR filtering*

All of the raw data were analyzed together using MaxQuant version 1.6.7.0. Spectra were searched against Arabidopsis TAIR10 genome, which was complemented with reverse decoy sequences and common contaminants by MaxQuant. Carbamidomethyl cysteine was set as a fixed modification while methionine oxidation and protein N-terminal acetylation were set as variable modifications. Digestion parameters were set to "specific" and "Trypsin/P;LysC". Up to two missed cleavages were allowed. A false discovery rate less than 0.01 at both the peptide spectral match and protein identification level was required. The "second peptide" option was used to identify co-fragmented peptides. The "match between runs" feature of MaxQuant was not utilized.

**Statistical analyses**

Statistical analyses on the protein abundance and ubiquitination data were performed using TMT-NEAT Analysis Pipeline version 1.3 (https://github.com/nmclark2/TMT-Analysis-Pipeline) (Clark et al., 2020). First, the MaxQuant output table was trimmed to only include the columns needed for statistical analysis and the columns were re-labeled using the provided sample information. Contaminants and reverse hits were removed at this stage. Prior to normalization, the two technical replicates for the abundance run were combined. Next, data were normalized using the sample loading normalization method (Plubell et al., 2017). Finally, statistical analysis was performed on the normalized values using PoissonSeq (Li et al., 2012). All resulting p- and q-values are retained and reported in Supplemental Tables 2 and 4 so that readers can use their preferred cutoff for selecting proteins and PTM sites that change after treatment for follow up studies. For our analyses, we classified protein groups and PTMs differentially accumulating as indicated in the Results and Discussion.

**Plasmid construction**

Full-length cDNA clones in pENTR/D-Topo were obtained from the Arabidopsis Biological Resource Center (ABRC) for the following transcription factors: CIB1/AT4G34530 (TOPO-U14-E03), CIL2/AT3G23690 (TOPO-U07-C08), and STOP1/AT1G34370 (TOPO-U16-

C08). Lysine sites of interest were mutated to arginine (K>R) using the Q5 Site-Directed Mutagenesis Kit (New England Biolabs, E0554) as described by the manufacturer. Primers for site directed mutagenesis were designed with NEBaseChanger and are listed in Supplemental Table 6. Positive clones with the desired mutations were confirmed by restriction enzyme digest and Sanger sequencing at the ISU DNA facility. Both WT and K>R pENTR/D-Topo clones were subsequently cloned into a modified p35S-LUC destination vector named “pDO18” using LR Clonase (Thermo Fisher Scientific). pDO18 was generated by amplifying the Firefly luciferase coding sequence from pBGL7.0 and inserted into the SpeI restriction sites in the pCMS44 vector, which is derived from p2BGW7. pCMS44 was created by mutating p2BGW7 using the Q5 site-directed mutagenesis kit and primers Q5SDM_7/16/2018_F and Q5SDM_7/16/2018_R to contain XhoI and KpnI restriction sites between the attR2 and T35S sequences. The wild-type (WT) and mutated K>R TF clones in the pDO18 backbone were verified by restriction digest and transformed into Agrobacterium strain GV3101.

### References

**Clark, N. M., Nolan, T. M., Wang, P., Song, G., Montes, C., Guo, H., Sozzani, R., Yin, Y., and Walley, J. W.** (2020). Integrated omics networks reveal the temporal signaling events of brassinosteroid response in Arabidopsis. *bioRxiv* Advance Access published September 5, 2020, doi:10.1101/2020.09.04.283788.

**Li, J., Witten, D. M., Johnstone, I. M., and Tibshirani, R.** (2012). Normalization, testing, and false discovery rate estimation for RNA-sequencing data. *Biostatistics* **13**:523–538.

**Plubell, D. L., Wilmarth, P. A., Zhao, Y., Fenton, A. M., Minnier, J., Reddy, A. P., Klimek, J., Yang, X., David, L. L., and Pamir, N.** (2017). Extended Multiplexing of Tandem Mass Tags (TMT) Labeling Reveals Age and High Fat Diet Specific Proteome Changes in Mouse Epididymal Adipose Tissue. *Molecular & Cellular Proteomics* **16**:873–890.

**Song, G., Hsu, P. Y., and Walley, J. W.** (2018). Assessment and Refinement of Sample Preparation Methods for Deep and Quantitative Plant Proteome Profiling. *PROTEOMICS* **18**:1800220.

**Song, G., Montes, C., and Walley, J. W.** (2020). Quantitative Profiling of Protein Abundance and Phosphorylation State in Plant Tissues Using Tandem Mass Tags. In *Plant Proteomics: Methods and Protocols* (ed. Jorrin-Novo, J. V.), Valledor, L.), Castillejo, M. A.), and Rey, M.-D.), pp. 147–156. New York, NY: Springer US.

Supplemental Table 1. Published studies reporting ubiquitin sites in Arabidopsis

| # of proteins with sites | # of reported sites | Site Identification Technique | Reference |
|---|---|---|---|
| 56 | 85 | K-e-GG footprint via LC-MS/MS | Maor et al. (2007) |
| 15 | 13 | K-e-GG footprint via LC-MS/MS | Saracco et al. (2009) |
| 109 | 120 | K-e-GG footprint via LC-MS/MS | Kim et al. (2013) |
| 1607 | 3009 | Lys-Gly label via LC-MS/MS | Walton et al. (2016) |
| 199 | 227 | K-e-GG footprint via LC-MS/MS | Aguilar-Herandez et al. (2017) |
| 3178 | 7130 | K-e-GG footprint via LC-MS/MS | This Work |

Several proteomic studies that identified ubiquitinated proteins but did not report site level information were not included (Manzano et al., 2008; Igawa et al., 2009; Svozil et al., 2014).

Supplemental Table 6. Primers used in this study.

| Primer Description | Primer sequence 5' - 3' | Clone info | Note |
|---|---|---|---|
| Q5SDM_4/10/2020_F | GAATTGGAGAGAACGGATTATATTC | AT4G34530 K166R | CIB1 |
| Q5SDM_4/10/2020_R | CTTCGTCACTTTAGATGAATC | AT4G34530 K166R | CIB1 |
| Q5SDM_4/10/2020_F | GACGATGCTAGGCCTCCTGAG | AT3G23690 K180R | CIL2 |
| Q5SDM_4/10/2020_R | CTTATTGTCGCCTTTACTATCG | AT3G23690 K180R | CIL2 |
| 5SDM_4/10/2020_F | GATGAGTACAGAACAGCAGCTG | AT1G34370 K271R | STOP1 |
| Q5SDM_4/10/2020_R | TCCATGCCCTCTCATATG | AT1G34370 K271R | STOP1 |
| Q5SDM_7/16/2018_F | cccgctcgagTCCCGCGGCCATGCTAGA | N/A | for creating pCMS44 plasmid |
| Q5SDM_7/16/2018_R | atatcggtaccTATCACCACTTTGTACAAGAAAGCTGAACG | N/A | for creating pCMS44 plasmid |

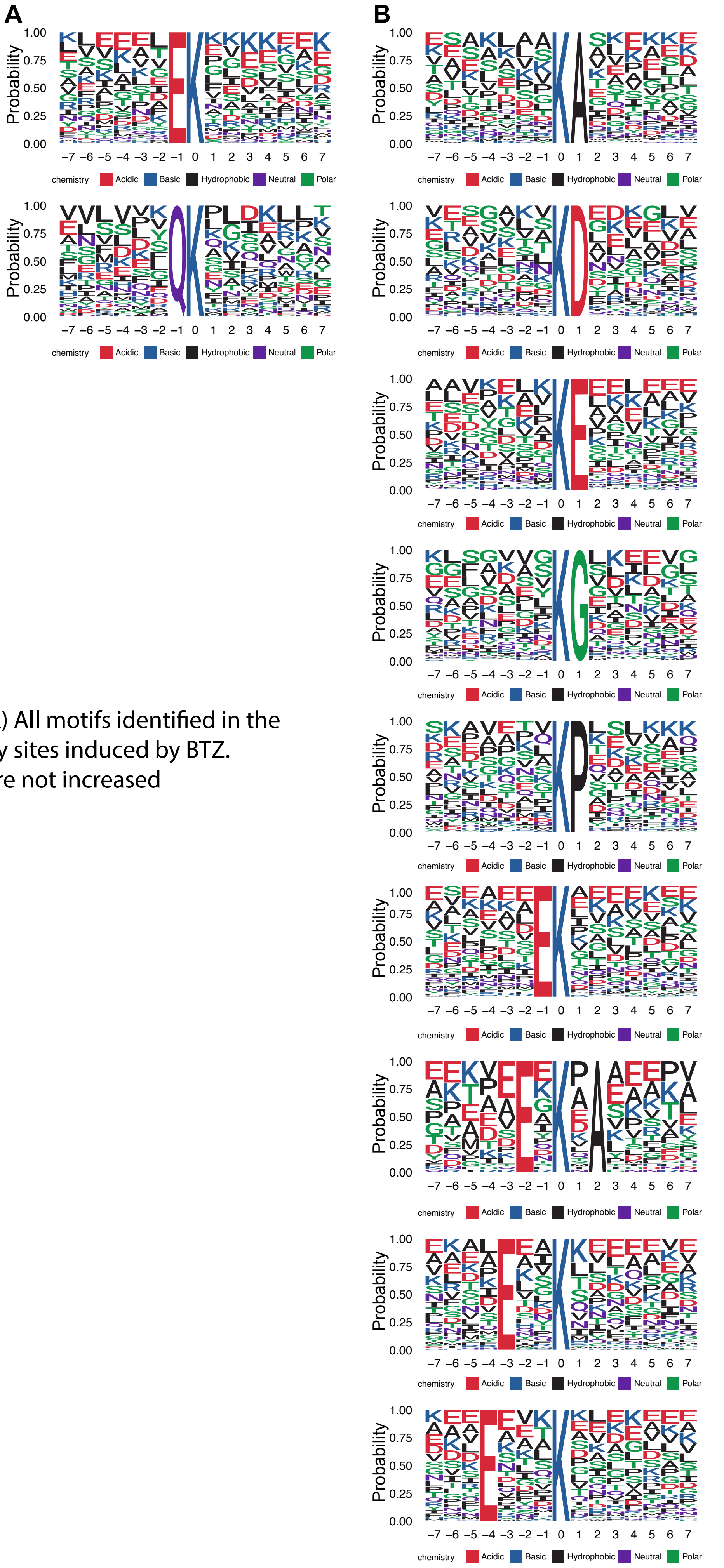


Supplemental Figure 1. Ubiquitin motif analysis. (A) All motifs identified in the 14 amino acid window surrounding the 1,336 diGly sites induced by BTZ. (B) Motifs identified among the diGly sites that were not increased following BTZ treatment.
